# Identification and Mechanistic Characterization of a Peptide Inhibitor of Glycogen Synthase Kinase (GSK3β) Derived from the Disrupted in Schizophrenia 1 (DISC1) Protein

**DOI:** 10.1101/2020.06.18.159665

**Authors:** Stephanie Saundh, Debasis Patnaik, Steve Gagné, Josh Bishop, Sean Lipsit, Samat Amat, Narsimha Pujari, Anand Krishnan Nambisan, Robert Bigsby, Mary Murphy, Li-Huei Tsai, Stephen Haggarty, Adelaine Kwun-Wai Leung

## Abstract

Glycogen Synthase Kinase 3-beta (GSK3β) is a critical regulator of several cellular pathways involved in neuroplasticity and is a potential target for neurotherapeutic development in the treatment of neuropsychiatric and neurodegenerative diseases. The majority of efforts to develop inhibitors of GSK3β have been focused on developing small molecule inhibitors that compete with ATP through direct interaction with the ATP binding site. This strategy has presented selectivity challenges due to the evolutionary conservation of this domain within the kinome. The Disrupted in Schizophrenia (DISC1) protein, has previously been shown to bind and inhibit GSK3β activity. Here, we report the characterization of a 44-mer peptide derived from human DISC1 (hDISCtide) that is sufficient to both bind and inhibit GSK3β in a non-competitive mode that is distinct from classical ATP competitive inhibitors. Based on multiple independent biochemical and biophysical assays, we propose that hDISCtide interacts at two distinct regions of GSK3β: an inhibitory region that partially overlaps with the binding site of FRATide, a well-known GSK3β binding peptide, and a specific binding region that is unique to hDISCtide. Taken together, our findings present a novel avenue for developing a peptide-based selective inhibitor of GSK3β.

## INTRODUCTION

Approximatley 16% of the world’s population suffers from neuropsychiatric disorders every year^1, 2^. While current treatments are efficacious, a significant proportion of patients either suffer from adverse side-effects or show decreased responsiveness to current treatments. Most major classes of neuropsychiatric drugs share similar mechanisms of action, which limits the therapeutic options for patients who do not respond well to the existing pharmaceuticals. In order to address this need, identifying novel molecular targets is a priority. One likely target is the multifunctional constitutively active Ser/Thr enzyme, glycogen synthase kinase 3-beta (GSK3β), which participates in the regulation of multiple cellular pathways including those responsible for cell proliferation, cell architecture, and neurodevelopment^3^. As a result of this diversity, a dysregulation of GSK3β has been implicated in various conditions, including mood disorders, Alzheimer’s disease, diabetes, and cancer^4^. More than 40 GSK3β substrates have been confirmed to date, and over 500 other potential substrates await validation^5, 6^. Clinical translation of GSK3β inhibitory molecules for neuropsychiatric diseases face three major challenges that include poor selectivity among closely related kinases and other CNS targets, blood-brain barrier (BBB) penetrability, and more significantly, chronic toxicity^7^. Although tremendous advances have been made in the development of highly selective GSK3β inhibitors^7, 8^, there still exists an opportunity to address novel mechanisms of inhibition that may overcome certain limitations, such as high cellular ATP levels and selectivity across the kinome.

As a basally active enzyme involved in a plethora of signaling pathways, GSK3β activity has multi-layered regulatory mechanisms that include post-translational modification, substrate priming, and protein complex formation. Regulation of the Wnt/β-catenin signaling pathway relies on partitioning GSK3β into distinct complexes. At rest, a subset of GSK3β is localized within a multi-protein unit called the destruction complex where the scaffold protein, Axin, binds to both GSK3β and its substrate β-catenin. GSK3β activity is enhanced ∼24,000 fold towards Axin-bound β-catenin compared to free β-catenin^9^. GSK3β acts to hyper-phosphorylate β-catenin which is unstable and is then targeted for degradation. In the presence of an external Wnt signal, Axin and GSK3β are sequestered into another protein complex called the signalosome where GSK3β activity is inhibited by the phosphorylated C-terminal tail of the Wnt co-receptor LRP5/6 occupying its substrate-binding site^10^. As a result, more stable β-catenin accumulates in the cytoplasm and translocates to activate Wnt-responsive genes in the nucleus. Both overabundant and insufficient activity of the Wnt/β-catenin pathway is linked to abnormal neurodevelopmental processes that contribute to the development of several psychiatric diseases^11^.

The psychiatric disease risk factor, Disrupted in Schizophrenia 1 (DISC1), is a scaffold-like protein that binds and inhibits GSK3β’s function in the Wnt/β-catenin pathway to modulate neuroprogenitor proliferation in the mouse model^12^. Moreover, DISC1’s inhibition of GSK3β seems to be specific to the Wnt/β-catenin pathway. In the same study, manipulating DISC1 expression had no effect on the inhibitory phosphorylation of Ser9 on GSK3β, which occurs in other signaling pathways involving GSK3β^12^. Additionally, not all GSK3β substrates are affected by the changes of DISC1 expression^12^. Therefore, DISC1-mediated inhibition of GSK3β offers an innovative approach in designing novel GSK3β inhibitors that are pathway and substrate selective. In the present study, we investigated the inhibition mechanism of a 44-residue peptide from human DISC1 (hDISCtide) that is sufficient to both interact with and inhibit GSK3β’s kinase activity. Our biochemical and biophysical data provide basis to an inhibition model whereby hDISCtide possesses two distinct regions that interact with GSK3β, one that acts as an anchor, and the other one which is necessary for inhibition.

## RESULTS AND DISCUSSION

Previously in an *in vitro* assay, a 44-residue region of the mouse DISC1 protein was shown to inhibit and bind to human GSK3β^12^. Here, we confirmed that the equivalent human sequence (hDISCtide), which is 70% identical, also inhibits human GSK3β in an *in vitro* kinase assay (Figure 1A). Using a phospho-primed glycogen synthase peptide (GSP2) as the substrate, hDISCtide inhibited recombinant GSK3β with an IC_50_ of 95 nM (Figure 1B). We also confirmed that hDISCtide could physically interact with GSK3β as assessed through a surface plasmon resonance (SPR) assay. In order to ensure a stable baseline, we generated recombinant N- and C-terminal double His-tagged GSK3β (His-GSK3β-His). Using this SPR assay, hDISCtide exhibited very slow binding kinetics to GSK3β with minimal dissociation after 10 minutes (Figure 1C). The surface of the SPR chip was regenerated after each injection so that fresh protein could be captured for each injection of hDISCtide. The sensorgrams at various hDISCtide concentrations were fitted to the OneToOne model (Equation 1, chi^2^ = 0.88), with a *k*_a_ of 495 (±31.4) M^-1^s^-1^ and *k*_d_ of 2.49×10^−4^ (±8.91×10^−6^) s^-1^. Due to the slow binding kinetics, the binding affinity cannot be estimated from the binding equilibrium dissociation constant (*K*_D_=*k*_a_/*k*_d_). Additionally, we were unable to obtain saturation binding at a practical concentration range to measure affinity by steady-state analysis. To cross reference the *K*_D_ value of hDISCtide with GSK3β we developed a microscale thermophoresis (MST) assay, which detects binding based on changes in molecular motions along a temperature gradient. Thus, MST differs from SPR in that binding is detected from free molecules in solution, whereas SPR requires one molecule to be immobilized. Using this MST assay the *K*_D_ of hDISCtide to GSK3β was measured to be 5.7 μM (Figure 1D).

**FIGURE 1.**
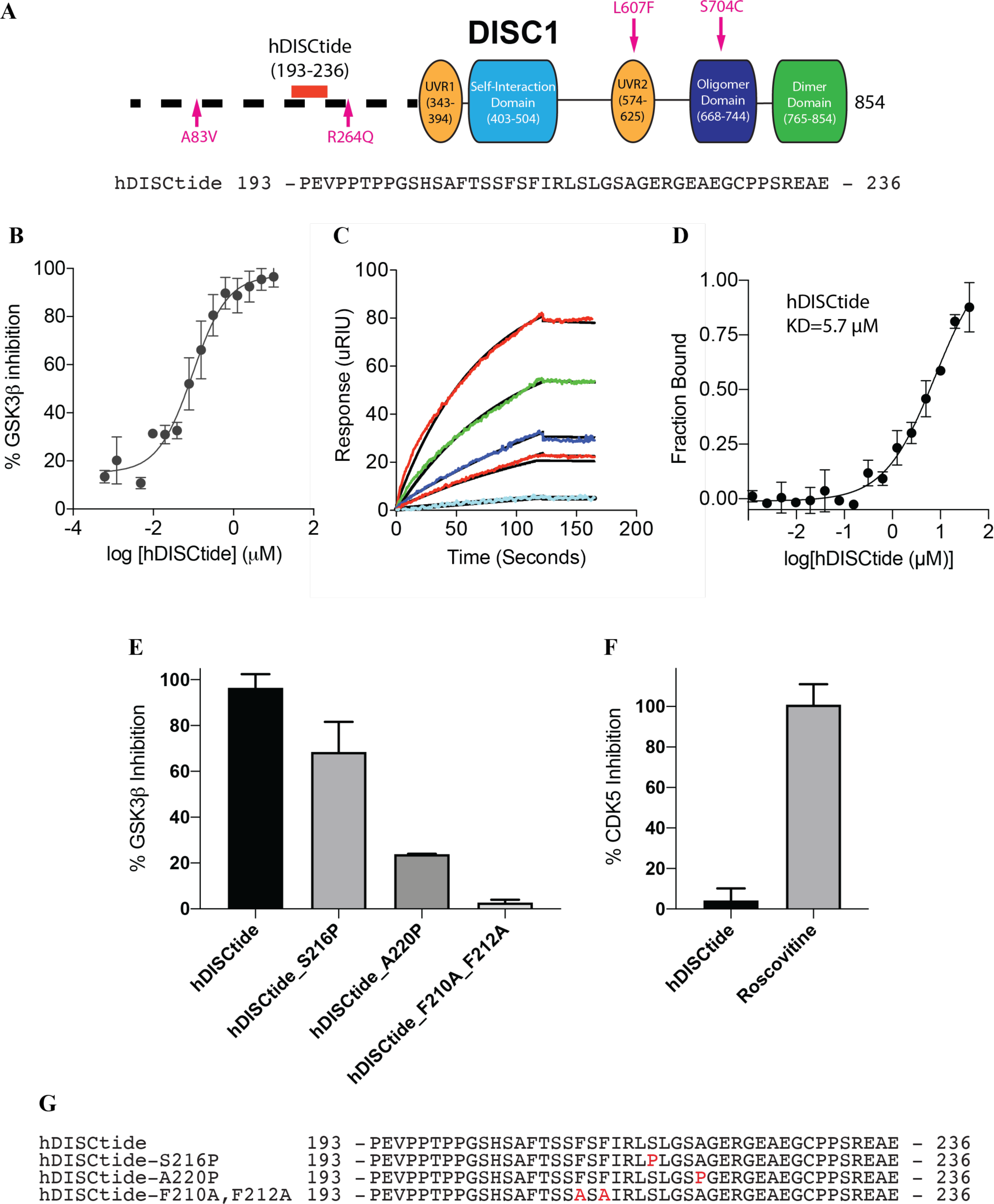
**A) Schematics of the full-length DISC1 isoform.** hDISCtide (residues 193-236) from the full-length protein is the most potent GSK3β inhibitory region of DISC1. The N-terminal region of DISC1 is predicted to be disordered, whereas the C-terminal domains contain several regions with high helical content that can self-associate or interact with other proteins^29^. Two stretches of the sequence are similar to the UVR-like domain, which is a conserved protein-protein interaction domain folded into an anti-parallel first identified in the bacterial endonuclease^30^. No other sequence similarity was noted in other regions of the protein^29^. A83V, R264Q, L607F, and S704C are disease-associated variants of DISC1. A83V, R264Q, and L607F have been shown to impact Wnt/GSK3β signaling while S704C does not have any effect^31^. **B) hDISCtide inhibition of GSK3β by ADP-Glo assay.** hDISCtide inhibits GSK3β with an IC_50_ of 95 nM. The reaction contains 10 nM GSK3β, 15 µM ATP, and 9 µM of the substrate peptide GSP2. Reaction was allowed to proceed for 20 min. The data were fitted with the equation for sigmoidal dose-response (variable slope) to estimate the IC_50_ (R^2^=0.9411). **C) hDISCtide binds GSK3β with slow kinetics.** ∼500-2000 RU of double His-tagged GSK3β was captured on a Ni-NTA chip. The concentration of hDISCtide injected was ranged from 1.79-28.6 µM. The surface was regenerated after every injection as little dissociation of the peptide was observed. The data were fitted to a OneToOne model with *k*_a_=495 M^-1^s^-1^ and *k*_d_=2.49×10^−4^ s^-1^ (Chi^2^=0.88). **D) Binding affinity of hDISCtide to GSK3β measured by microscale thermophoresis assay.** Fraction of GSK3β bound with hDISCtide. The data were fitted with the equation for sigmoidal dose-response (variable slope) to estimate the *K*_D_ of 5.7 µM (R^2^=9720). **E) hDISCtide inhibition is sequence specific.** Percent GSK3β inhibition by 10 µM of hDISCtide and its mutants. Single site mutation of S216P reduces the inhibition potency while A220P shows no inhibition. The double mutant F210A_F212A completely abolished hDISCtide inhibition. **F) hDISCtide does not inhibit CDK5, a structurally similar kinase to GSK3β.** Roscovitine, a small molecule general kinase inhibitor, is the positive control of the CDK5 assay. Percent CDK5 inhibition by 11 µM of hDISCtide and roscovitine. **G) Sequences of hDISCtide for Figure E.** Top sequence is the wild-type sequence. Highlighted in red are the mutated residues.

To determine the sequence specificity of the inhibition and binding to GSK3β, we next examined a peptide composed of the same amino acids as hDISCtide but synthesized in a scrambled order. This peptide did not show any efficacy as an inhibitor of GSK3β (data not shown). Subsequently, several mutagenesis analyses were performed to evaluate the impact of different amino acid substitutions on GSK3β inhibition. A single-site mutation of A220P completely abolished hDISCtide inhibition, whereas S216P reduced inhibition potency (Figure 1E, G). A peptide with a double mutation of F210A and F212A was unable to inhibit GSK3β (Figure 1E, G).

To determine whether hDISCtide acts as a general kinase inhibitor, its effect on a cyclin dependent kinase 5 (CDK5), a close homolog of GSK3β, was examined. CDK5 shows 24.5% identity and 38.1% similarity in amino acids compared to GSK3β. Most reported GSK3β inhibitors that are not selective also affect CDK5 activity^13^. To assess how selective hDISCtide is to GSK3β, we used an ADP-Glo assay to test the inhibition potency of hDISCtide against CDK5 in a reaction containing ATP (at approximate K_m_ values) and the CDK5 substrate peptide (PKTPKKAKKL) derived from histone H1. For comparison, we included an assay containing the small molecule non-selective inhibitor of GSK3β, roscovitine. hDISCtide did not show any inhibition against CDK5, whereas, roscovitine showed potent inhibition (Figure 1F). Thus, hDISCtide appears to posess selectivity against GSK3β, and therefore the inhibition mechanism of hDISCtide is likely distinct from that of a general kinase inhibitor.

Simple inhibition modality can be described based on the affinity of the inhibitor to different enzyme states along the reaction coordinate of the enzyme. The inhibitor can bind exclusively to the free enzyme (E) (competitive inhibition), exclusively to the enzyme-substrate complex (ES) (uncompetitive inhibition), or to both species (noncompetitive inhibition). In noncompetitive inhibition, the degree to which the inhibitor changes the binding affinity of the substrate to the enzyme is described by the constant α. When α is significantly larger than 1, the inhibitor completely excludes the binding of the substrate. When α=1, the inhibitor has the same affinity as the free enzyme and ES complex. When α>1, the inhibitor binds with a higher affinity to the free enzyme. When α<1, the inhibitor binds with a higher affinity to the ES complex^14^.

To investigate the inhibition modality of hDISCtide against GSK3β, we evaluated how different concentrations of hDISCtide influence the V_max_ and K_m_ of the two substrates (ATP and GSP2) used in our kinase assays. To determine if hDISCtide can compete for ATP binding, GSK3β enzymatic reactions were carried out with varying hDISCtide and ATP concentrations in the absence of GSP2 using a time-resolved fluorescence energy transfer (TR-FRET) kinase assay (Supplementary Figure 1A). Similarly, to determine if hDISCtide can compete directly with the substrate peptide GSP2, GSK3β enzymatic reactions were carried out with varying hDISCtide and GSP2 concentrations while the ATP concentration was fixed at the approximate K_m_ value of 15 µM using the ADP-Glo assay (Supplementary Figure 1B). Both datasets were fitted to the Michaelis-Menten equation, followed by an estimation of the K_m_ and V_max_ for the respective substrate at each hDISCtide concentration. As a control, we also performed a similar kinetic analysis with CHIR-99021, a well-established, ATP competitive, small molecule inhibitor of GSK3β, using the TR-FRET assay with varying hDISCtide and ATP concentration in the absence of GSP2 (Supplementary Figure 1C). First, pure competitive inhibitors should have no effect on V_max_^14^, as expected and observed for CHIR-99021 (Figure 2C-i). Since ^ATP^V_max_ and ^GSP2^V_max_ both decreased curvilinearly with increasing hDISCtide concentration, we can conclude that hDISCtide does not directly compete with ATP or GSP2 substrate (Figure 2A-i, 2B-i)^14^. Second, uncompetitive inhibitors should have no effect on the ratio V_max_/K_m_^14^. Since this ratio decreases curvilinearly for the three datasets, we can rule out uncompetitive inhibition (Figure 2A-iii, 2B-iii, 2C-iii)^14^. Finally, the α constant was deduced from the K_m_ replots. ^ATP^K_m_ increased curvilinearly with increasing hDISCtide concentration, indicating non-competitive inhibition with α>1^14^ (Figure 2A-ii). In comparison, the linearity of the ^ATP^K_m_ replot for CHIR-99021 is a signature trend for a pure ATP competitive inhibitor (Figure 2C-ii). Conversely, ^GSP2^K_m_ decreased curvilinearly with increasing hDISCtide concentration, indicating non-competitive inhibition with α<1^14^ (Figure 2B-ii). In summary, hDISCtide is a non-competitive inhibitor relative to both ATP and GSP2. Based on the α constants, hDISCtide has a higher affinity for free GSK3β compared to ATP-bound GSK3β. In contrast, hDISCtide has a higher affinity for the GSP2-bound GSK3β compared to an enzyme where the substrate peptide binding pocket is unoccupied.

**FIGURE 2.**
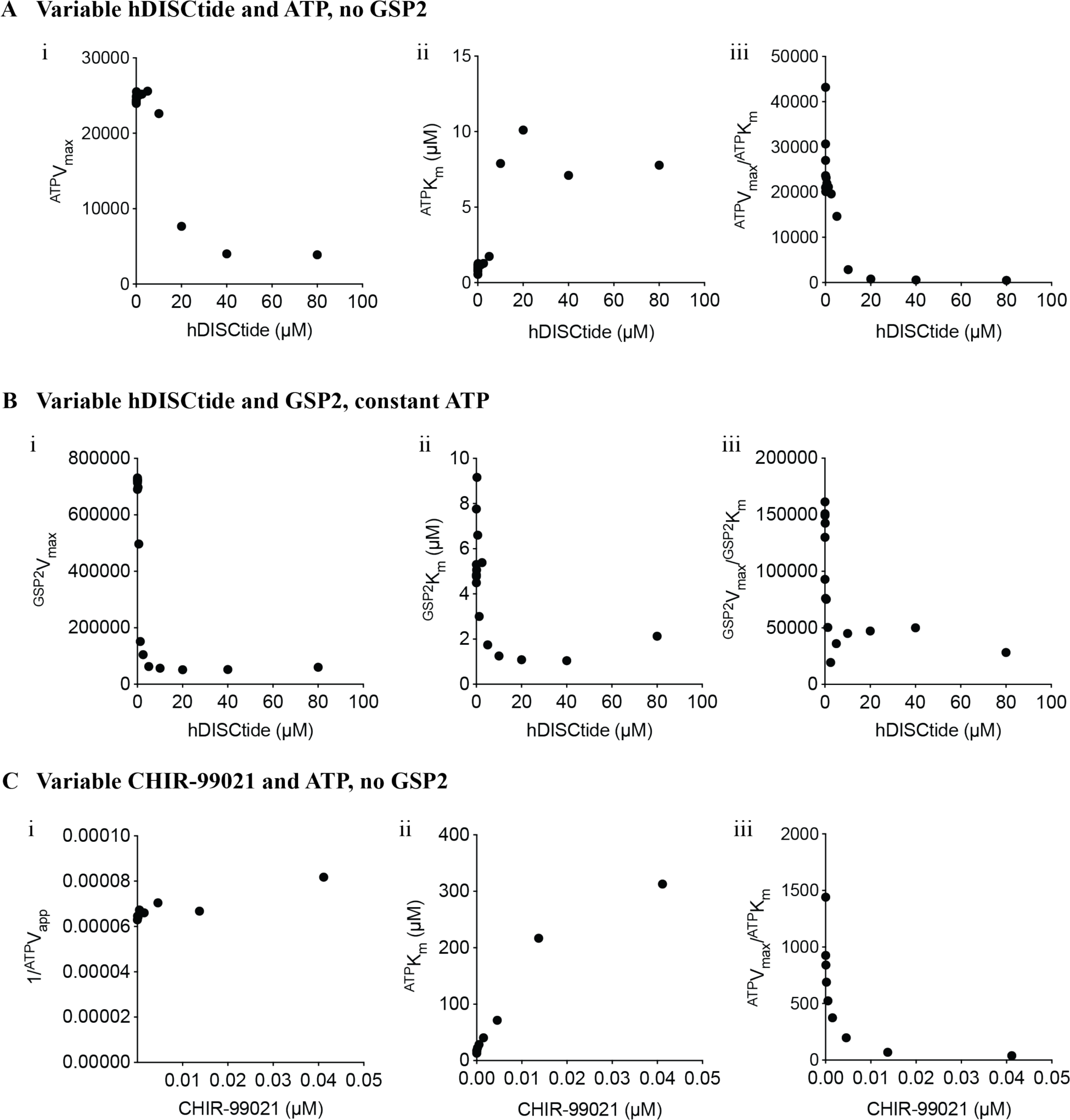
Replots of the Michaelis-Menten kinetic paremeters show that hDISCtide is a non-competitive inhibitor of GSK3β with respect to ATP and GSP2. **A)** To determine if hDISCtide can compete directly with ATP, GSK3β enzymatic reactions were carried out in the absence of GSP2. ^ATP^V_max_ and ^ATP^K_m_ were obtained from data fitted to the Michaelis-Menten equation (Supplementary Figure 1A). **i)** Replots of ^ATP^V_max_ as a function of increasing hDISCtide concentration. ^ATP^V_max_ decreases curvilinearly with increasing hDISCtide concentration, indicating an absence of competition^14^. **ii)** Replots of ^ATP^K_m_ as a function of increasing hDISCtide concentration. ^ATP^K_m_ appears to increase curvilinearly with increasing hDISCtide concentration, leading us to conclude that the mechanism of inhibition is non-competitive with α>1^14^. **iii)** Replots of ^ATP^V_max_/^ATP^K_m_ showing the ratio decreases curvilinearly with increasing hDISCtide concentration, thus ruling out un-competitive inhibition^14^. **B)** To determine if hDISCtide can compete directly with the substrate peptide GSP2, GSK3β enzymatic reactions were carried out with a fixed concentration of ATP and varying concentrations of hDISCtide and GSP2. ^GSP2^V_max_ and ^GSP2^K_m_ were obtained from data fitted to the Michaelis-Menten equation (Supplementary Figure 1B). **i)** Replots of ^GSP2^V_max_ as a function of increasing hDISCtide concentration. ^GSP2^V_max_ decreases curvilinearly with increasing hDISCtide concentration, indicating an absence of competition^14^. **ii)** Replots of ^GSP2^K_m_ as a function of increasing hDISCtide concentration. ^GSP2^K_m_ appears to decrease curvilinearly with increasing hDISCtide concentration, leading us to conclude that the mechanism of inhibition is non-competitive with α<1^14^. **iii)** Replots of ^GSP2^V_max_/^GSP2^K_m_ showing the ratio decreases curvilinearly with increasing hDISCtide concentration, thus ruling out un-competitive inhibition^14^. **C)** As a comparison, similar mechanistic analyses were performed with the well-established ATP competitive small molecule inhibitor of GSK3β, CHIR-99021. GSK3β enzymatic reactions were carried out in the absence of GSP2. ^ATP^V_max_ and ^ATP^K_m_ were obtained from data fitted to the Michaelis-Menten equation (Supplementary Figure 1C). **i)** Replots of 1/^ATP^V_app_ as a function of increasing hDISCtide concentration. Varying hDISCtide concentration has no effect on 1/^ATP^V_app_, indicating pure competitive inhibition^14^. **ii)** Replots of ^ATP^K_m_ as a function of increasing CHIR-99021 concentration. ^ATP^K_m_ increases linearly with increasing CHIR-99021 concentration, a signature trend for ATP competitive inhibitors. **iii)** Replots of ^ATP^V_max_/^ATP^K_m_ showing the ratio decreases curvilinearly with increasing hDISCtide concentration, thus ruling out un-competitive inhibition^14^.

Since hDISCtide is a non-competitive inhibitor relative to both ATP and the substrate peptide GSP2, the binding site of hDISCtide likely involves regions outside of the ATP and substrate binding pockets. The C-terminal domain of GSK3β contains a protein binding site where two well-characterized peptides (AxinGID and FRATide) bind^9, 15^. Both of these peptides are derived from scaffold proteins that localize subsets of cellular GSK3β to different compartments of a cell. The crystal structures of GSK3β/AxinGID and GSK3β/FRATide show that the peptides partially share the same binding site at the C-terminal domain of GSK3β (Figure 5A-B, highlighted in blue)^9, 15^. AxinGID binds GSK3β as a single helix (Figure 5A), whereas FRATide binds GSK3β as two α-helices bent at a 90 degree angle (Figure 5B)^9, 15^. The plasticity of the peptide-binding loop can accommodate the two unique conformations of AxinGID and FRATide. To test the hypothesis that hDISCtide shares binding site with AxinGID or FRATide at a similar location on GSK3β, we sought to compare their binding kinetic parameters with that of hDISCtide measured by SPR. First, we confirmed the binding of either AxinGID or FRATide to GSK3β using a Biolayer Interferometry (BLI) assay, which follows similar biophysical assay principles compared to SPR. BLI yielded a *K*_D_ of 3.6 μM and 63.6 μM for FRAtide and AxinGID respectively (Figure 3A, Supplementary Figure 2A). The low affinity of AxinGID was likely due to the inclusion of only residues that were observed to be ordered in the crystal structure^9^. Removing the flexible flanking residues appear to affect its affinity.

**FIGURE 3.**
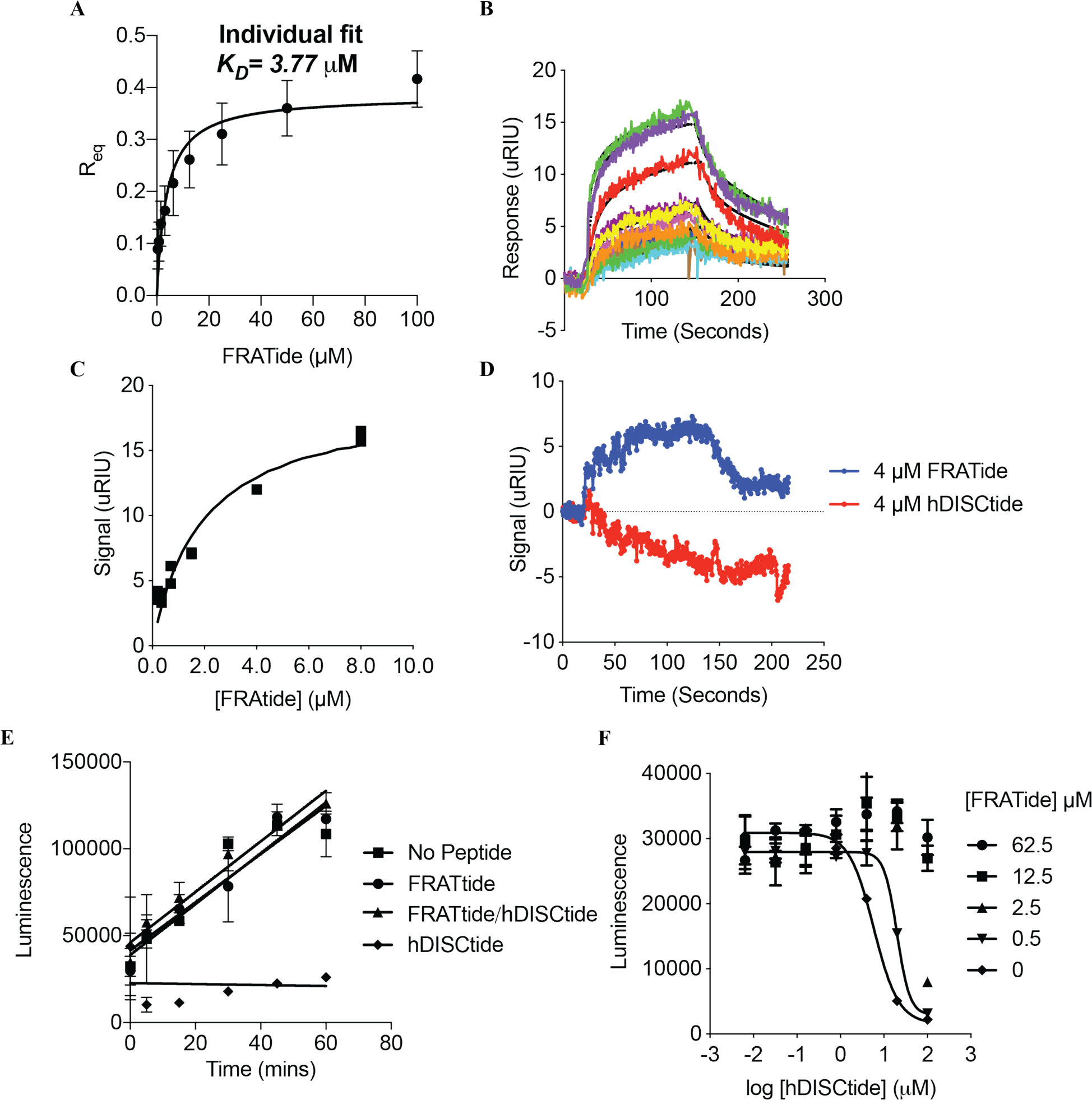
FRATide and hDISCtide partially shares binding site. **A)** FRATide binds GSK3β with a *K*_D_ of 3.8 µM measured by the BLI assay. The equilibrium dissociation constant (*K*_D_) was estimated by steady-state analysis using the response signal at equilibrium for each FRATide concentration (Supplementary Figure 3). The data were fitted globally with the 1:1 binding model with the equation Response=(R_max_*Conc.)/*K*_D_ + Conc), from which the *K*_D_ is estimated at the concentration where 50% of R_max_ is reached. **B-C)** Real-time binding analysis of FRATide to GSK3β by SPR. FRATide binding to GSK3β has faster kinetics compared to hDISCtide. ∼500-2000 RU of double his-tagged GSK3β was captured on a NiNTA chip. The concentrations of FRATide injected were 0.2, 0.35, 0.7, 1.5, 4, 8 µM. The data was fitted to a OneToOne TwoState model with *k*_a1_=9940 M^-1^s^-1^, *k*_d1_=6.50×10^−2^ s^-1^, *k*_a2_=1.62×10^−2^ M^-1^s^-1^, *k*_d2_=8.09×10^−3^ s^-1^ (Chi^2^=0.44), *K*_D_=3.2 µM. The *K*_D_ from equilibrium analysis is 1.9 µM. **D) Real-time analysis of the co-binding of hDISCtide and FRAtide to GSK3β by SPR.** 3 µg/ml (∼64 nM) double his-tagged GSK3β was incubated with 4 µM of hDISCtide before capturing (∼553 RU). Injecting 4 μM of hDISCtide did not generate any response (red). Subsequent injection of 4 μM FRATide gave a response with comparable kinetics as that observed on a surface captured with unbound GSK3β in B (blue). **E-F) FRATide abolishes hDISCtide inhibitory activity against GSK3β**. E) ADP-Glo luminescent assay to compare GSK3β activity in the presence of FRATide and/or hDISCtide. 25 µM of FRATide and/or hDISCtide was added to each condition containing 25 nM of GSK3β, 9 µM GSP2, 100 µM ATP in reaction buffer. F) Double titration curves of hDISCtide and FRATide. FRATide at a concentration of 2.5 µM and higher eliminated inhibtion potency of hDISCtide against GSK3β. At 0.5 µM concentration, FRATide increased the IC_50_ of hDISCtide ∼3 fold.

**FIGURE 4.**
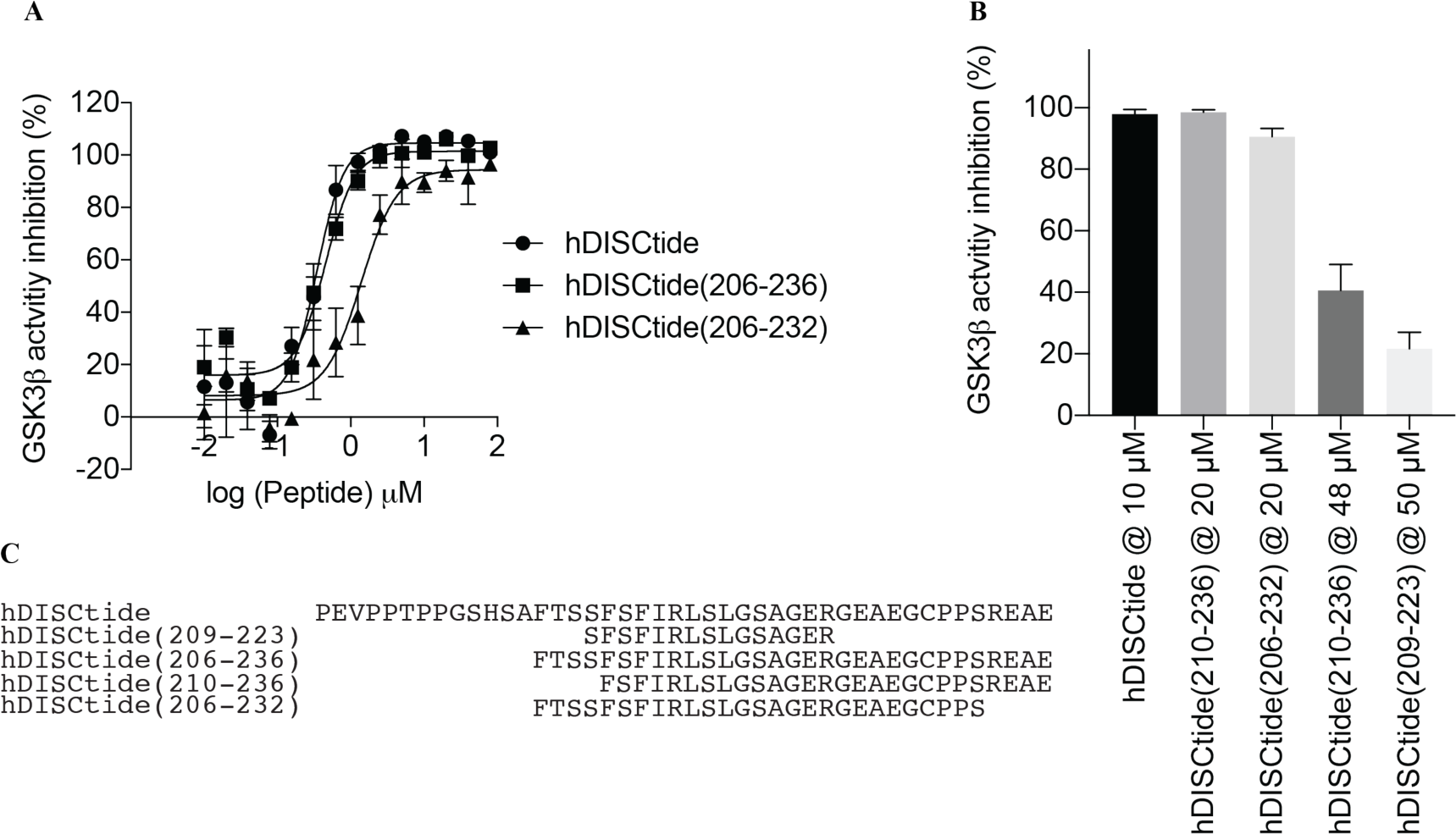
Inhibitory action of hDISCtide truncations on GSK3β activity. Different truncations of hDISCtide were tested for their ability to inhibit GSK3β using the ADP-Glo assay. The N-terminal 13 residues were dispensible to hDISCtide inhibition potency whereas trimming the C-terminus by 4 residues were detrimental to the inhibitory action of hDISCtide. **A)** Dose-response inhibition of hDISCtide and its truncated constructs against GSK3β. **B)** Percent inhibition of GSK3β by the different hDISCtide truncation constructs. **C)** Sequences of the hDISCtide constructs for panels 5A and 5B.

**FIGURE 5.**
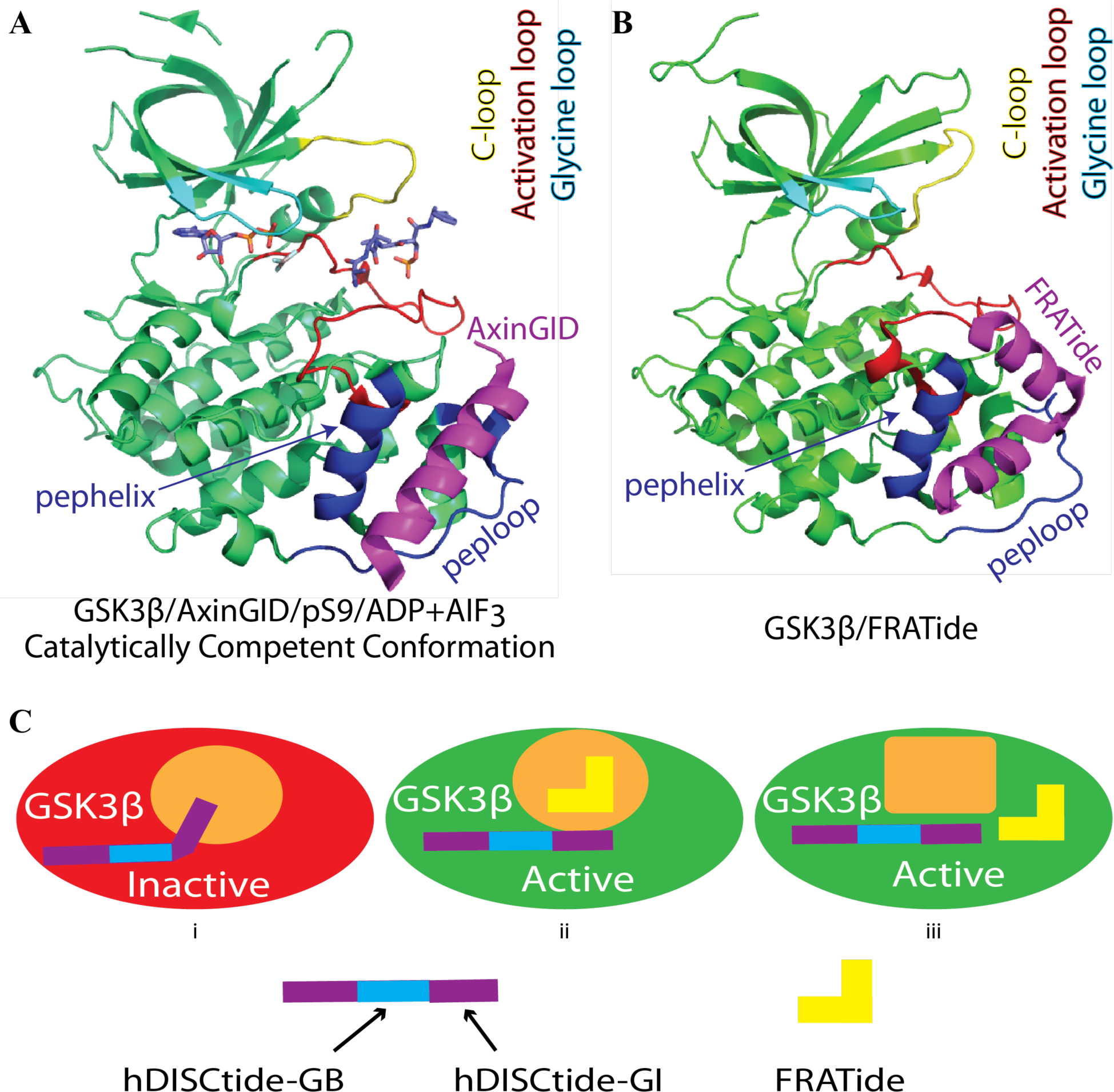
Crystal structures of GSK3β in different conformations. **A)** Catalytically competent conformation of GSK3β. GSK3β bound to the scaffold peptide Axin (magenta). The catalytic conformation of the enzyme consists of the glycine-rich loop (cyan), the C-loop (yellow), and the activation loop (red). The scaffold binding regions consist of a helix and a loop (blue). The substrate peptide binding pocket is bound by the pseudo-substrate peptide formed by the N-terminus (residues 6-10) with a phosphorylated Ser9. The pseudo substrate peptide, ADP, and the transition analog AIF3 are shown in stick model (PDB:4NU1). **B)** GSK3β bound to the scaffold peptide FRATide (magenta) (PDB:1GNG). **C)** Models of hDISCtide inhibition mechanism. GSK3β activity depends on the three regions forming a catalytic conformation (Figure 6A), represented by the orange rectangle. hDISCtide could inhibit activity by preventing the formation of a catalytic competent GSK3β, as represented by an orange oval. hDISCtide has two binding regions: hDISCtide-GB and hDISCtide-GI. hDISCtide-GI binds to the orange oval to block GSK3β from forming catalytic competent conformation (i.e. orange rectangle). FRATide only affects the binding of hDISCtide-GI but not hDISCtide-GB. It can either block hDISCtide-GI from interacting to its binding site or changes the conformation of the GI binding site such that hDISCtide-GI can no longer bind.

Since FRATide showed higher affinity binding to GSK3β in the BLI assay, we sought next to perform a comparative analysis of the binding kinetics of FRATide and hDISCtide to GSK3β using the SPR-based assay. In contrast to the slow binding kinetics observed for hDISCtide, FRATide bound GSK3β with a fast on- and off-rate. Unlike hDISCtide, which never reached saturation in our experimental setting, the binding sites for FRATIde were saturated within two minutes of injection (Figure 3B). We were able to measure the *K*_D_ by steady-state and kinetic analysis (Figure 3B). The affinity of FRATide to GSK3β was determined to be 1.9 µM by steady-state analysis (Figure 3C). The kinetic data did not fit the standard OneToOne model, but instead fitted best to a OneToOne TwoState model (Equation 2). The OneToOne TwoState model describes the presence of an initial weaker interaction event, followed by a subsequent change to a stronger interaction. The initial weaker interaction event could be contributed by GSK3β binding to only one of the two helices of FRATide, followed by the stronger interaction formed with the additional contacts from the second FRATide helix. Nevertheless, the affinity obtained from this kinetic model was 3.2 μM, which is comparable to that obtained from steady-state analysis (1.9 μM).

Taking advantage of the substantial difference in GSK3β binding kinetics between hDISCtide and FRATide that could be measured with the SPR assay, we next asked whether FRATide can compete with hDISCtide for binding at a similar site. Since hDISCtide binding kinetics were slow, we first incubated GSK3β with 4 μM of hDISCtide for 20 mins in solution before capturing the GSK3β-hDISCtide complex on the sensor chip. After immobilization, subsequent injection of hDISCtide (4 μM) showed no detectable binding, confirming that the binding sites for hDISCtide on GSK3β had been saturated (Figure 3D – red). In contrast, FRATide was still able to interact with the GSK3β-hDISCtide complex, and presented similar binding kinetics (Figure 3D – blue) when compared to FRATtide interacting with GSK3β alone (Figure 3B). This observation indicates that FRATide can still access its binding site on hDISCtide-bound GSK3β, suggesting that the FRATide and hDISCtide binding sites do not overlap. To further test the notion that FRATide and hDISCtide bind at distinct sites on GSK3β, we investigated how FRATide influences hDISCtide inhibition of GSK3β. Since FRATide binding to GSK3β blocks its activity toward non-phospho-primed substrates (e.g. Axin), but does not affect GSK3β’s activity toward phospho-primed subsrtratres like that of glycogen synthase^16^, we reasoned that FRATtide should not affect the inhibition potency of hDISCtide in our GSK3β activity assay using a glycogen synthase-derived peptide substrate. We first confirmed that FRATide on its own does not inhibit GSK3β (Supplementary Figure 2B). However, contrary to our expectation, the inhibition of GSK3β by hDISCtide was abolished in the presence of FRATide (Figure 3E). To explore this effect more, a double titration experiment was performed. In the presence of 0.5 μM FRATide, the IC_50_ of hDISCtide increased ∼3-fold. At concentrations of 2.5 μM and above, FRATide completely abolished the inhibition of hDISCtide even at 100 μM (Figure 3F). Thus, these data show that the binding of FRATide blocks the ability of hDISCtide to form an inhibitory interaction with GSK3β.

The contradictory interpretation from the SPR and ADP-Glo assays might be explained by considering the size of the peptides. Given that the size of hDISCtide (44 a.a.) is twice as long as that of FRATide (25 a.a.), the protein-protein-interface (PPI) of GSK3β/hDISCtide could be larger than that of GSK3β/FRATide particularly if hDISCtide binds in an extended conformation. FRATide may disrupt the region of the PPI of GSK3β/hDISCtide that is essential for its inhibitory function while other regions of hDISCtide remain bound to GSK3β. In this proposed model, there are two distinct functional regions of hDISCtide: one that acts as an anchor and forms specific interactions with GSK3β and one that is required for its inhibitory function. Our previously published studies are consistent with this model. When overlapping 15-mer peptide segments spanning the mouse DISCtide region were tested for GSK3β binding, only residues from 211 to 225 showed GSK3β binding by SPR; however, these truncated peptides did not inhibit GSK3β kinase activity^12^. In our hands, we also observed that the equivalent human 15-mer, hDISCtide(209-225), only bound, but did not inhibit human GSK3β (data not shown). This indicates that the regions flanking the central 15-mer are essential for GSK3β inhibition. To identify which end of the central 15-mer was responsible for hDISCtide inhibitory activity, several truncated hDISCtide peptides were evaluated using the ADP-Glo assay. From the N-terminus, 13 residues were dispensable to the inhibition potency, whereas trimming the C-terminus by just four residues greatly reduced the inhibition potency (Figure 4A-C). Based on this data, we propose that hDISCtide(209-225) comprises the anchor that interacts with a unique site on GSK3β (Figure 5C –”GB”), whereas hDISCtide(226-236) is the region (Figure 5C – “GI”) responsible for GSK3β kinase inhibition. The presence of FRATide interferes with the interactions between hDISCtide-GI and GSK3β (Figure 5C).

The crystal structure of GSK3β trapped in a catalytically competent conformation (PDB:4NU1) shows that three regions are important for catalysis: 1) the glycine loop (cyan), which changes conformation upon substrate peptide binding; 2) the C-loop (yellow), which provides key specificity to substrate peptides and engages in conformational change upon substrate peptide binding; 3) the activation loop (red), which forms the substrate peptide binding surface opposite to the C-loop. (Figure 5A)^10^. Since our kinetic data show that hDISCtide inhibition is non-competitive with ATP or the substrate peptide, hDISCtide could inhibit GSK3β by preventing the above three regions of GSK3β from forming a catalytic competent conformation (Figure 5C-i). hDISCtide inhibition can only occur if hDISCtide-GI can interact with its binding site on GSK3β (Figure 5C-i orange oval). The method by which FRATide overcomes hDISCtide’s inhibition of GSK3β could occur in one of two ways: 1) FRATide could physically block the hDISCtide inhibitory binding site (Figure 5C-ii); or 2) FRATide binding could induce a conformational change in the hDISCtide-GI binding site (Figure 5C-iii). The crystal structure of GSK3β/FRATide shows that FRATide binds at the C-terminal domain of GSK3β clamped by a helix (pepHelix: 262-273) and a loop (pepLoop: 285-299) (Figure 5B, segment coloured in blue). These regions can potentially interact directly or indirectly with hDISCtide-GI while hDISCtide-GB interacts with regions outside of this FRATide binding site.

Collectively, this work provides new insights on how GSK3β is regulated through its association with DISC1. Further investigation is required to understand the inhibition mechanism of DISC1 against GSK3β within a relevant cell type in the context of the full-length protein. DISC1 regions outside of the 44 residue DISCtide could interact with other proteins and influence which subcellular GSK3β pool that can be inhibited by DISC1. For example, GSK3β could exist in the destruction complex in the absence of Wnt signal, the signalosome in the presence of Wnt signal, or remain free in the cytoplasm. Addressing which pool of GSK3β DISC1 could interact and inhibit will shed light on the temporal involvement of DISC1 in the regulation of Wnt signaling – does DISC1 also modulate the basal level of GSK3β activity or only during Wnt activation? Moreover, it would be important to consider other DISC1 interactors that have been found to regulate Wnt-GSK3β/β-catenin signaling and modulate neural progenitor proliferation during early embryonic cortical development, such as Dixdc1^17^. Dixdc1 binds at the C-terminus (596-852) of DISC1 and has a calpain homology domain that interacts with actin. It would be interesting to determine if Dixdc1 influences DISC1’s inhibition of GSK3β that occurs at the N-terminus. Some short DISC1 isoforms that contain the GSK3β inhibitory region, but lack the Dixdc1 binding region, have been reported to be upregulated in the hippocampus of schizophrenic patients^18^. If Dixdc1 affects DISC1’s inhibition of GSK3β, different ratios of long and short DISC1 isoform will influence the overall Wnt/β-catenin signaling and the pathophysiology of schizophrenia.

As the minimal GSK3β inhibitory region of DISC1, hDISCtide has therapeutic potential for rescuing the loss of GSK3β inhibitory function upon loss of DISC1. Moreover, since the binding site of hDISCtide partially overlaps with that of AxinGID and FRATide, hDISCtide can potenitally disrupt GSK3β association with their respective scaffold proteins, Axin and FRAT. AxinGID is derived from Axin that is a key component in the Wnt/β-catenin signaling pathway. This work suggests that hDISCtide can potentially normalize Wnt signaling by inhibiting Axin-dependent GSK3β phosphorylation of β-catenin. FRATide is a fragment from FRAT, which was initially thought to segregate GSK3β binding away from Axin when the Wnt/β-catenin pathway is activated^16^. However, FRAT was subsequently found to be non-essential for the Wnt/β-catenin pathway in higher organisms^19^. Instead, the main function of FRAT appears to be a shuttle that carries nuclear GSK3β to the cytoplasm when the PI3K/AKT/mTORC1 signaling pathway is activated^20, 21^. By changing the subcellular localization, FRAT can control GSK3β activity towards cytoplasmic or nuclear substrates in response to PI3K/AKT/mTORC1 signalling. More recently, high expression of FRAT has been reported to confer rapamycin resistance in several cancer cell lines by reducing the population of nuclear GSK3β that catalyzes inhibitory phosphorylation of nuclear proteins that repress cell growth^21^. Therefore, it is tempting to argue that hDISCtide could also normalize the PI3K/AKT/mTORC1 signaling pathway by preventing FRAT-dependent shuttling of GSK3β out of the nucleus and promotes GSK3β phosphorylation of its nuclear substrates.

GSK3β has attracted substantial pharmaceutical investments due to its implication in multiple diseases including type 2 diabetes, Alzheimer’s disease (AD), bipolar disorder, and cancer. A recent paper outlining AstraZeneca’s 20+ years of experience in developing small molecules that target GSK3β in hope to treat AD showed that work on their best clinical candidates were terminated due to toxicity^22^. Our work presents a novel therapeutic strategy in designing peptide-based inhibitors of GSK3β. Our data indicates that hDISCtide is a non-competitive inhibitor to both ATP and the substrate peptide GSP2. In particular, inhibitors that compete with the substrate peptide binding site offer new guidance for drug design that are more specific for GSK3β^23^. However, peptide-based CNS therapeutics face their own challenges since they have short half-lives and a low penetrability of the BBB. Intensive research is currently underway to develop new strategies that will allow therapeutic peptides to be specifically transported to brain tissues. An example of this is peptide modification (N-terminal acylation, glycosylation, halogenation) used to increase BBB perviousness and peptide delivery through the use of nanoparticles which have shown recent success^24, 25^. Future structural and cell-based studies will be necessary to evaluate the efficacy and further develop hDISCtide as a therapeutic for CNS disorders.

## METHODS

### Peptides

All peptides used in this study were synthesized by NeoBioLab or Genscript. FRATide: DPHRLLQQLVLSGNLIKEAVRRLHS (GenScript); AxinGID: VEPQKFAEELIHRLEAVQ (Genscript); hDISCtides sequences are as listed in Figures 1A, G, 4C.

### Small Molecule Kinase Inhibitors

CHIR-99021 was purchased from Stemgent (04-0004-10). Roscovitine was purchased from Calbiochem (186692-46-6).

### GSK3β Cloning, Expression, and Purification

Four different sources of GSK3β was used in this study: 1) GST-GSK3β purchased from BP-Biosciences (BPS Biosciences Cat 40007); 2) GSK3β made in house for the competition experiment between hDISCtide and FRATide by ADP-Glo assays; 3) His-GSK3β-His made in house for the binding assays using SPR; 4) 6X-HisGSK3β purchased from Signalchem (G09-10H-05) for BLI and MST experiments. *Cloning:* The codon optimized human GSK3β gene was synthesized and subcloned into the pST44 backbone containing an N-terminal histidine tag (pST44-HisNGSK3β)^26^. An additional histidine tag was inserted at the C-terminus of the reading frame (pST44-HisNGSK3βHis).

#### Protein Overexpression and Purification

All constructs were transformed in BL21(DE3)star. Seed cultures were inoculated in 1-2 L TB media. The cultures were induced with 0.15 mM IPTG at OD=0.1-0.2 at 16-17 °C for 35-40 hours. Cells were resuspended in lysis buffer (50 mM Hepes pH 7.2, 50 mM NaCl, 5% Glycerol, 1 mM PMSF) and lysed by cell disruption (Constant systems LTD 0.75KW, two passes with 25K Psi) at 4 °C. The lysate was centrifuged at 40,000 x g for 40-60 mins. The supernatant was diluted 1:10 in dilution buffer (50 mM Hepes pH 7.2, 5% glycerol, 50 mM NaCl) and loaded on an SP-Sepharose-FF column equilibrated with Buffer A (50 mM Hepes pH 7.2) and washed for 20-40 column volume. Bound proteins were eluted in a linear gradient with 100% Buffer B (Buffer A + 1 M NaCl). Fractions containing GSK3β were pooled and loaded onto a NiNTA column equilibrated with Buffer C (50 mM Tris pH 7.5, 300 mM NaCl, 12.5 mM imidazole) and washed for 10-20 column volumes. His-GSK3β was eluted in a 100% step gradient with Buffer D (Buffer C + 250 mM imidazole). Protein was pooled and cleaved with TEV at 1:10 and dialyzed in 50 mM Hepes pH 7.2, 500 mM NaCl, 5 mM beta-mercaptoethanol overnight. For some batches, cleaved GSK3β was already pure enough. Otherwise, an additional size-exclusion chromatography was performed with buffer (50 mM Hepes pH 7.2, 500 mM NaCl, 2 mM MgCl2, 1 mM TCEP). Pure GSK3β was flash-frozen in liquid nitrogen and stored at -80 °C. For SPR studies, His-GSK3β-His protein was stored without TEV cleavage.

### Surface Plasmon Resonance (SPR)

SPR studies were conducted using the Reichert 2-channel system (Model: 2SPR). The Nickel Nitrilotriacetic Acid (Ni-NTA) Sensor Chip (Reichert CA# 13206065) was first prepared by flowing a NiSO_4_ solution overtop to capture the Ni^+^ ions which primed it for protein capture. SPR assay was conducted in running buffer (50 mM Tris pH 7.5, 150 mM NaCl, 5 mM MgCl_2_, 0.01% Tween 20, 1 mM TCEP). Between 500-2000 RU of His-GSK3β-His were captured onto the Ni-coated surface. Once the baseline was stable (<5 RU/min), analytes were injected at 25 µl/min for 2 mins association time and 2 mins dissociation time. The NiNTA chip surface was regenerated with the stripping buffer (20 mM NaPhos pH 8.5, 300 mM NaCl, 50 mM EDTA) or the regeneration buffer (50 mM Hepes pH 7.2, 300 mM NaCl, 250 mM imidazole) and 50 mM NaOH solution. Each regeneration solution was injected at 100 µl/min for 2 mins. Each chip could be regenerated for up to ∼10 times before the capture level decreased significantly. Results were analyzed and fitted using TraceDrawer 1.6.1 (Ridgeview Instruments AB). For datasets that required regeneration after every injection, local fitting of Bmax for each curve was performed to account for differences in capture level.

### ADP-Glo Kinase Assay

Reagents for the ADP-Glo assay were purchased from Promega (Cat V9102). For GSK3β kinase assay, reactions were performed with 9 μM GSK3β substrate peptide (YRRAAVPPSPSLSRHSSPHQ S(PO_3_H_2_)EDEEE), 15 μM ATP, 10-25 nM GSK3β in reaction buffer (50 mM Tris pH 7.5, 5 mM MgCl_2_, 0.01% Brij-35, 3 mM DTT). The protein, substrate, and GSK3β ligands were preincubated for 15 minutes before the addition of ATP. Reactions were allowed to proceed for 30 minutes. ADP production was monitored by luminescence as described in the Promega manual.

For CDK5 kinase assay, the reaction contains 10 nM CDK5-p25 (a gift from Kosik lab), 40 µM ATP, 20 µM cdk5 substrate peptide (PKTPKKAKKL, Anaspec #60026-1). The reaction was allowed to proceed for 25 minutes. The luminescence due to kinase activity was quantified using a luminescence plate reader as per the manufacturer’s recommendations (Envision).

### Time-Resolved Fluorescence Energy Transfer (TR-FRET) Assay

Reagents for TR-FRET assays were purchased from Perkin Elmer. Reactions were performed with 50 nM ULight-Glycogen Synthase (Ser641/pSer657) peptide (Perkin Elmer #TRF0131-M), 15 μM ATP, 1 nM GSK3β in the same reaction buffer as above. The protein, substrate, and GSK3β ligands were preincubated for 15 minutes before the addition of ATP. Reactions were allowed to proceed for 25 minutes. Reaction termination and detection were performed using 1X LANCE detection buffer with 20 mM EDTA and 2 nM LANCE Ultra Europium anti-phospho Glycogen Synthase (Ser641) (Perkin Elmer #TRF0220-M).

### Microscale Thermophoresis (MST) and Bioilayer Interferometry (BLI) Assays

The MST assays were performed as described previously^27^. Similarly, the BLI assays were conducted as described^28^, with the use of biotinylated 6X-His-GSK3β as the recombinant protein.

### Data Analysis

All data were analyzed using GraphPad Prism 8.0.1 (GraphPad Software Inc.).

## ABBREVIATIONS

ADP-GLO: Adenosine Di Phosphate Glow
ADP: Adenosine Di Phosphate
BBB: Blood Brain Barrier
CNS: Central Nerous System
DISC1: Disrupted in Schizophrenia 1
FRAT: Frequently Rearranged in Advanced T Cell Lymphoma
GS: Glycogen Synthase
GSK3β: Glycogen Synthase Kinase 3 Beta
GST: Glutathione S-transferase
NiNTA: Nickel Nitrilotriacetic Acid
SP-FF: SP Sepharose Fast Flow
SP-HP: SP Sepharose High Performance
SPR: Surface Plasmon Resonance
TEV: Tobacco Etch Virus
Wnt: Wingless/Integrated

## ACKNOWLEDGMENTS

We would like to thank Drs. David Sanders and David Palmer for critically reading the manuscript. Drs. Michal Boniecki and Jason Maley for help with the SPR instrument. We wish to thank Dr. Kelly L. Arnett and Harvard’s Center for Macromolecular Interactions for advice regarding Microscale Thermophoresis (MST). We want to thank Dr. Kenneth S. Sosik, UC Santa Barbara, for providing the recombinant CDK5/p25 enzyme preparation.

^**\***^This work was supported by a NIH grant (L.H.T; S.J.H.), a JELF grant from Canada Foundation for Innovation (Funding Reference No. JELF#31655) (A.K.W.L), an Establishment Grant from the Saskatchewan Health Research Foundation (A.K.W.L), a Discovery grant from Natural Sciences and Engineering Research Council of Canada (Funding Reference No. RGPIN-2014-06012) (A.K.W.L), internal grants from the Western College of Veterinary Medicine and the College of Medicine at the University of Saskatchewan (A.K.W.L), and the Stuart & Suzanne Steele MGH Research Scholars Program (S.J.H).

