## Supplemental Data for "Identification and Mechanistic Characterization of a Peptide Inhibitor of Glycogen Synthase Kinase (GSK3β) Derived from the Disrupted in Schizophrenia 1 (DISC1) Protein"

A

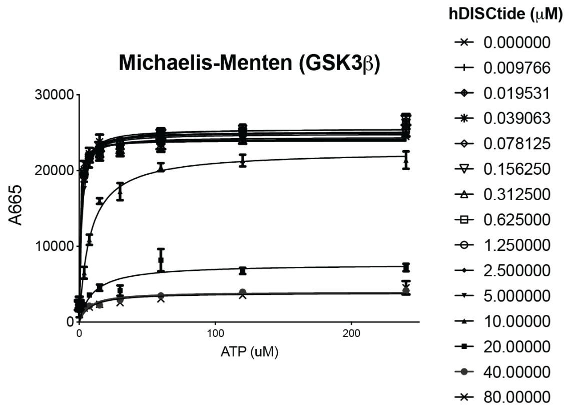


B

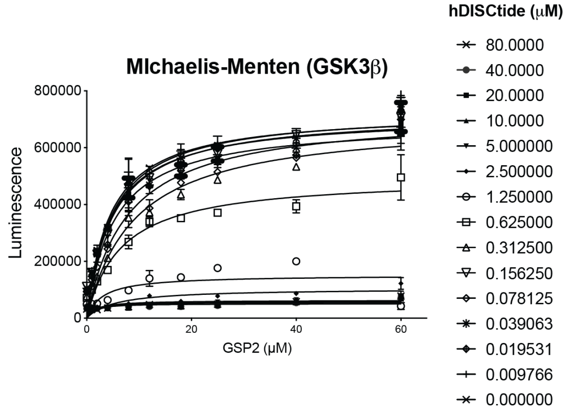


C

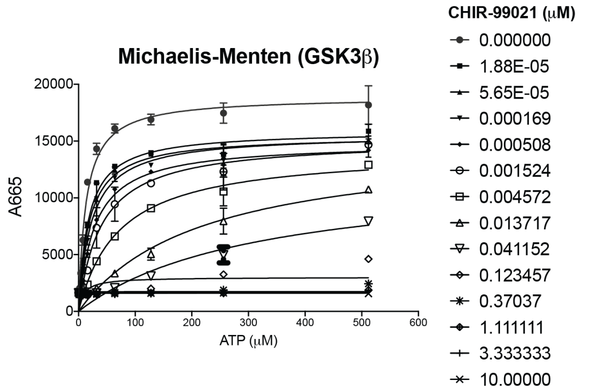


**SUPPLEMENTARY FIGURE 1**. **A)** Michaelis-Menten fitted curves of GSK3β enzymatic reactions with varying concentrations of hDISCtide and ATP in the absence of the substrate peptide GSP2, as measured in the TR-FRET assay. **B)** Michaelis-Menten fitted curves of GSK3β enzymatic reactions with varying concentrations of hDISCtide and GSP2 and a fixed concentration of ATP, as measured in the ADP-Glo assay. **C)** Michaelis-Menten fitted curves of GSK3β enzymatic reactions with varying concentrations of CHIR-99021 and ATP in the absence of the substrate peptide GSP2, as measured in the TR-FRET assay.

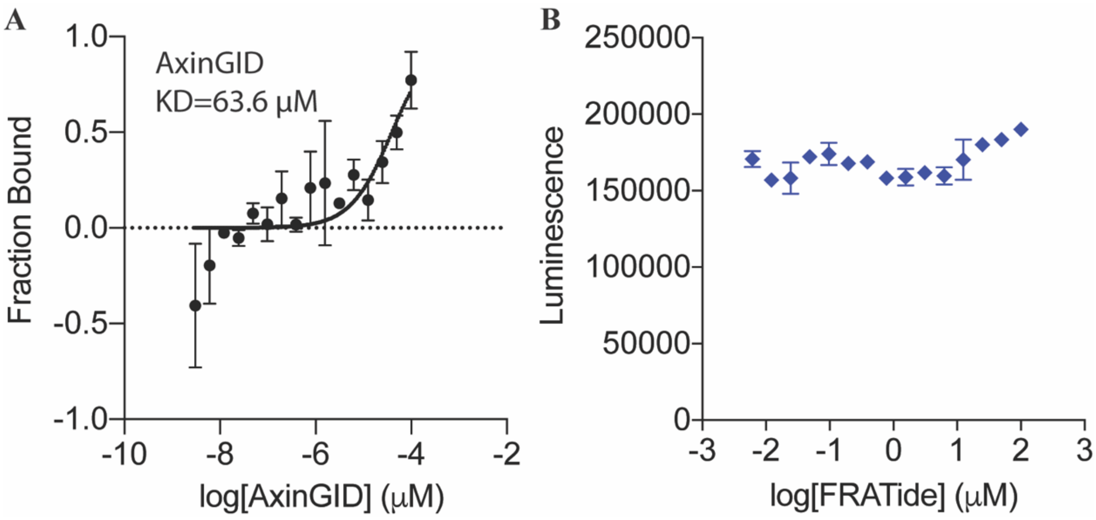


**SUPPLEMENTARY FIGURE 2**. **A)** AxinGID binds GSK3β with a $K_D$ of 63.6 µM measured by the BLI assay. **B)** FRATide does not inhibit the kinase activity of GSK3β in the ADP-Glo assay with GSP2 substrate peptide.

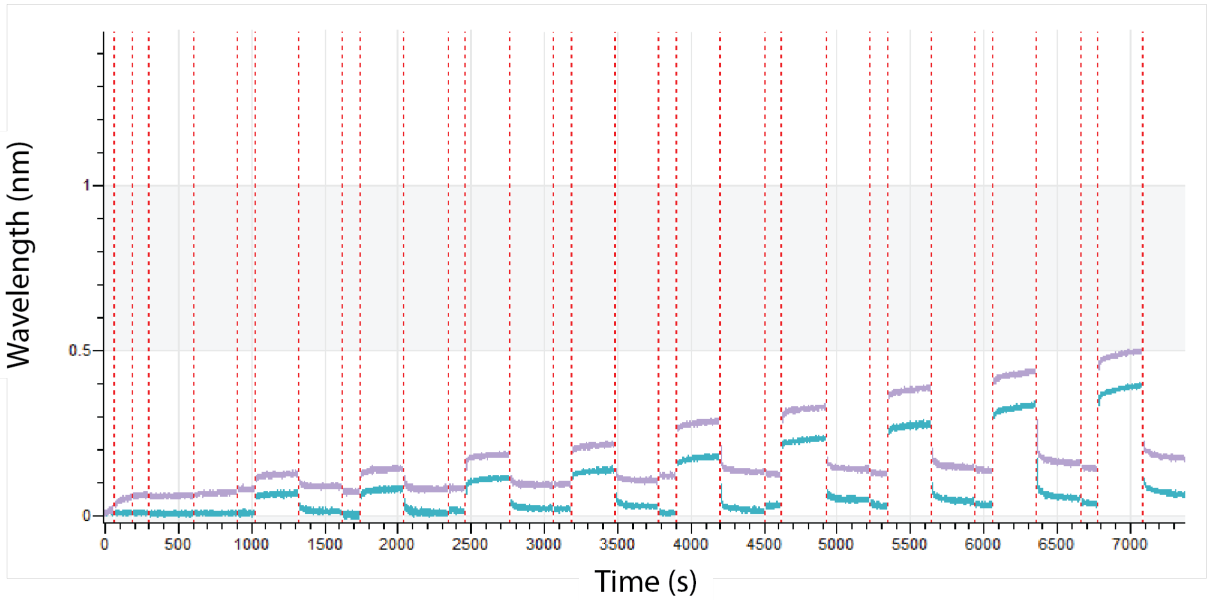


**SUPPLEMENTARY FIGURE 3**. **Biolayer interferometry-based biophysical assays of hDISCtide binding to GSK3β.** The reference subtracted real-time binding kinetics of FRAtide to GSK3β. A spectral shift is indicative of a binding interaction.

Equation 1: OneToOne model[1]

$$\frac{dY}{dt} = (k_a * c - k_d) * Y;$$

Y(t=0) = 0, signal from saturated target = Bmax

Y = recorded SPR signal

c = concentration of ligand

$k_a$ [1/(M*s)] = The association rate constant

$k_d$ [1/s] = The dissociation rate constant

Equation 2: OneToOne TwoState model[1]

$$\frac{dB}{dt} = -(k_{a1} * B * c \; - \; k_{d1} * A1B)$$

$$\frac{dA1B}{dt} = (k_{a1} * B * c \; - \; k_{d1} * A1B) \; - \; (k_{a2} * AB \; - \; k_{d2} * A2B)$$

$$\frac{dA2B}{dt} = (k_{a2} * A1B - k_{d2} * A2B)$$

$$Y = A1B \; + \; A2B$$

$$A1B(t = 0) = A2B(t = 0) = 0, B(t = 0) \; = \; Bmax$$

Y = recorded SPR signal

A1B = initial weaker complex

A2B = final stronger complex

B = unbound target

c = concentration of ligand

Bmax = maximum signal

$k_{a1}$ [1/(M*s)] = The association rate constant for the first binding state

$k_{d1}$ [1/s] = The dissociation rate constant for the first binding state

$k_{a2}$ [1/s] = The formation rate constant for the second binding state

$k_{d2}$ [1/s] = The disruption rate constant for the second binding state
